# Gaussian Process Inference Reveals Non-separability of Position and Velocity Tuning in Grid Cells

**DOI:** 10.64898/2026.02.02.703269

**Authors:** Linnie J. Warton, Surya Ganguli, Lisa M. Giocomo

## Abstract

Grid cells in medial entorhinal cortex (MEC) support spatial navigation by responding to multiple variables, including position, speed, and head direction. While tuning curves for each of these variables have been examined individually at the level of single-cells, less is known about the conjunctive coding of grid cells for these properties. To investigate this, we analyzed neural recordings of freely foraging rats and constructed four-dimensional (4D) tuning curves across 2D position and 2D velocity. In order to combat the sparse sampling of such a large behavioral space, we applied Gaussian Process (GP) methods to estimate firing rates at un-sampled points. Comparing GP model-derived tuning curves to those predicted by a fully separable model revealed that some cells exhibited significant non-separability of position and velocity tuning, and suggested a data coverage threshold necessary to observe this non-separability. In summary, our use of GPs allowed us to distinguish interactions in position-velocity tuning across a 4D behavioral space that have not been apparent in 2D analyses.

## Introduction

Animal survival depends on accurate spatial coding, which enables animals to determine their position in space and navigate to food, home, or a mate. Over the last several decades, many of the building blocks of an internal neural navigation system have been described in the mammalian medial entorhinal cortex (MEC). This system depends, in part, on MEC grid cells, which form periodic and hexagonal firing patterns that tile space and can provide a metric representation of distance traveled (Fyhn et al., 2004; Hafting et al., 2005). Grid cells can also exhibit conjunctive tuning for both position and other navigationally relevant variables, such as head direction (Sargolini et al., 2006), and can alter their firing in response to changes in running speed (Kropff et al., 2015; Hardcastle et al., 2017; Low et al., 2021). Several studies have shown that there are cells that code for multiple variables simultaneously, including grid cells in MEC, and that these cells can code for angular and linear velocity conjunctively as well (Hardcastle et al. 2017; Spalla et al. 2022). Furthermore, while grid cells were initially proposed to maintain a rigid metric across space, recent work indicates that their activity is also modulated by factors such as task demands (Boccara et al., 2019; Butler et al., 2019). However, the algorithms grid cells use to combine these multiple navigation variables remain incompletely understood.

Computational models such as continuous attractor networks (CANs) have proposed that grid cells form a toroidal manifold, a low-dimensional neural representation capable of maintaining position estimates through recurrent activity (McNaughton et al., 2006; Guanella et al., 2007; Burak and Fiete, 2009). Recent large-scale recordings in MEC have confirmed this structure, revealing toroidal manifolds within co-recorded modules of grid cells that differ in grid scale (i.e. the size of grid firing fields and the distance between these fields) and orientation (Stensola et al., 2012; Gardner et al., 2022). One such dataset, collected by Gardner et al. (2022), offers extensive recordings of grid cell modules in freely foraging rats and provides a unique opportunity to examine how spatial tuning varies across behavioral conditions.

Here, we leverage this data set to ask whether speed acts purely as a gain modulator of a neuron’s position encoding or whether speed and position interact in a more complex and non-separable way. Traditional methods for examining this question marginalize over one variable to examine tuning in another (e.g. assessing 2-dimensional [2D] position tuning marginalized over all velocities). In the simplest case, a behaviorally invariant spatial firing pattern would result in a stable grid cell firing pattern across all movement patterns, with speed modulating only the magnitude of firing. This would be mathematically equivalent to a factorizable, or separable, code for position and velocity. However, for grid cells, it remains unknown whether grid cells exhibit truly separable tuning for position and velocity or whether position and velocity are integrated to form a neural code across a higher-dimensional behavioral space.

A major challenge in addressing this question is obtaining comprehensive coverage across all combinations of position, speed, and directions. To address this challenge, we applied Gaussian Processes (GP) models, which use observed data points to infer mean estimates of unobserved points. GP models thus offer a principled approach to estimating neural activity from sparsely sampled data and have previously been used to estimate 2D grid cell tuning curves (Rule et al., 2023) and neural manifolds across speeds (Ye and Wessel, 2024). Here, we used GP models to visualize and quantify four-dimensional (4D) tuning curves of grid cells over position and velocity. We then compared these GP-estimated 4D tuning curves to those generated by a separable model in which speed and position tuning were independent. We found that in sessions with greater coverage of the 4D behavioral space, the GP model outperformed the separable model in predicting held-out neural activity. This finding points to both the behaviorally dependent nature of grid cell coding and the importance of high-dimensional models for understanding the neural representation of external space.

## Results

### Analyzing grid cell activity in an open field arena

We used a comprehensive dataset collected in ‘Toroidal topology of population activity in grid cells’ from Gardner et al., 2022. Briefly, rats were chronically implanted with Neuropixels silicon probes and neurons from the medial entorhinal cortex (MEC) were recorded as rats foraged for randomly scattered food in an open field (OF) arena (1.5 m by 1.5 meters) (Figure 1A). The data set consists of neural activity recorded from three rats: Q, R, and S. Rat R was recorded for two sessions over two days and Q and S were each recorded for one session (average session length = 64 minutes). Grid cells in MEC can be categorized into modules of cells that share the same grid field spacing, or scale. Cells within a module differ in terms of where grid fields are located (i.e. grid cell phase) but not the average distance between grid fields (Stensola et al., 2012). Since the dataset consists of only grid cells, our analyses considered cells within a given module separately. Gardner et al., reported three modules in rat R, two modules in rat Q, and one module in rat S.

**Figure 1:**
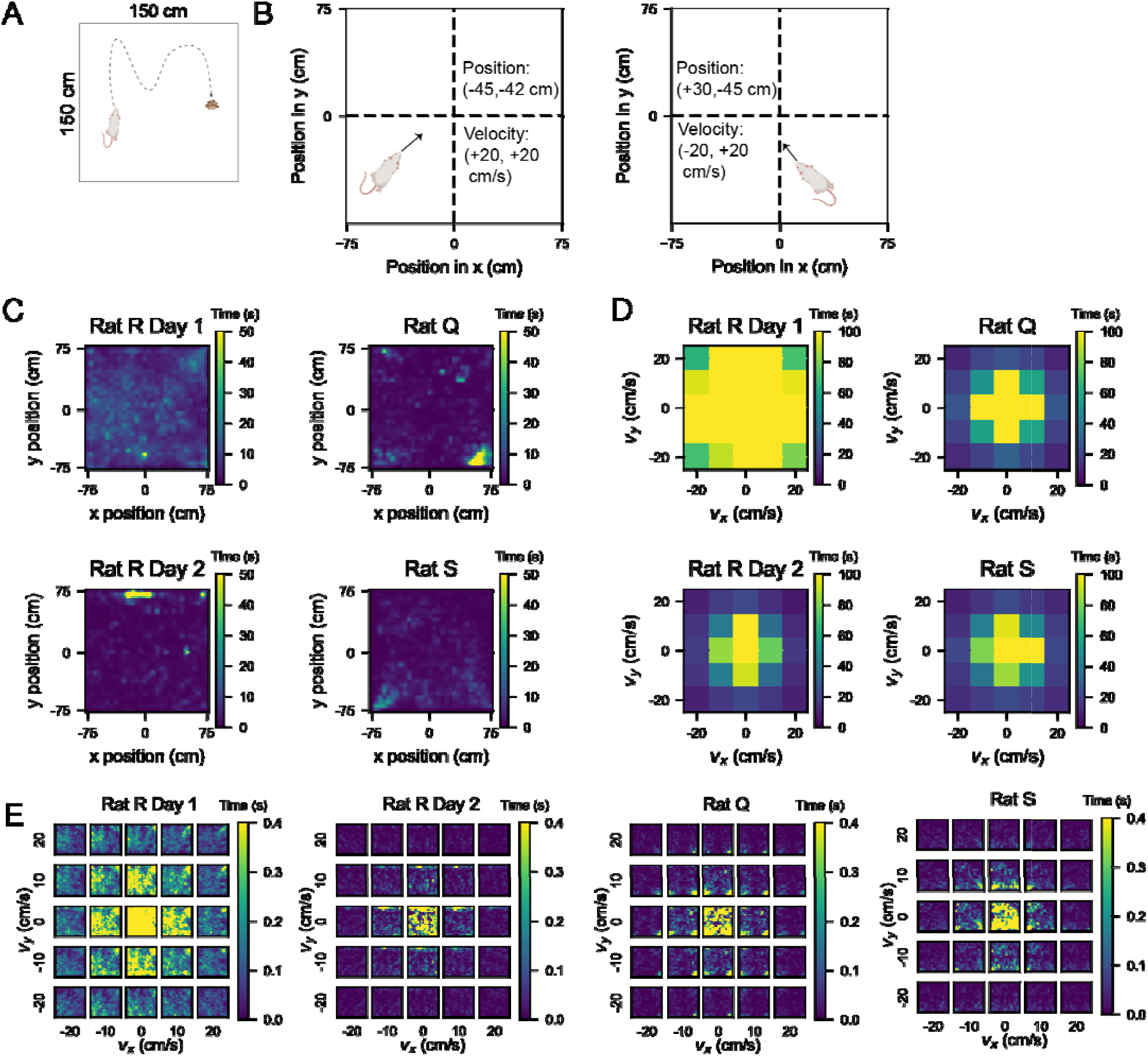
Coverage of the 4D conjunctive position-velocity space was lower than behavioral coverage of the 2D position and 2D velocity space independently. A) Schematic of the open field (OF) foraging task from Gardner et al. 2022, in which rats (R, Q and S) foraged for randomly scattered food rewards (yellow dots) in a 1.5 x 1.5 meter arena. B) Schematic illustrating velocity and position quantification. We denoted position as a 2D vector with the first element being the x-coordinate and the second the y-coordinate. Locations to the left of the center of the arena were denoted with negative x-coordinates (and to the right with positive x-coordinates), and locations below the center were denoted with negative y-coordinates (and above with positive y-coordinates). We represented velocity as a 2D vector to indicate speed and direction. Trajectories toward the left side of the arena featured a negative x-coordinate for velocity, whereas trajectories toward the right had a positive x velocity. Trajectories downward had a negative y velocity, and upward had positive y velocity.1 C) 2D histograms of position occupancy in the arena for each behavioral session. The color bar indicates the amount of time spent at each position bin, color coded for maximum (yellow) and minimum (blue) values. While there were a few restricted over-sampled positions, all position bins were sampled relatively evenly throughout each session. D) 2D histograms of occupancy in velocity space for each behavioral session. We limited our analysis to velocity within the range of −25 to +25 cm/s in both x and y coordinates, as the occupancy declined after these values. Plots color coded for maximum (yellow) and minimum (blue) values (amount of time spent in each velocity bin), with bins = 10 cm/s. E) Taking the 2D position (panel C) and 2D velocity (panel D) occupancy altogether, we visualized the time spent across bins in 4D position-velocity space. Each small square in the 5 x 5 heatmaps is a histogram at a particular velocity bin of time points spent in each position bin in the arena, as in (C). The outer edges of these 5 x 5 heatmaps, which represent the highest speed, show suboptimal coverage of 2D position in the arena. Each plot is color coded for maximum (yellow) and minimum (blue) values (seconds spent in each velocity bin).

Gardner et al. constructed 2D tuning curves by segmenting the arena into 3 cm x 3 cm bins and smoothing them with a Gaussian kernel. We recapitulated these tuning curves with 5 cm x 5 cm bins and observed the cardinal 60 degree symmetry of grid cell firing patterns in open field environments (Hafting et al., 2005). For the purposes of our analyses, we plotted position tuning curves with the x position ranging from −75 to +75 cm left to right and the y position ranging from −75 to +75 cm bottom to top (Figure 1B).

We overlaid a coordinate plane onto the open field (OF) arena such that the center of the arena corresponds to an x and y position of 0. Negative x positions extended to the left of the arena, and positive x positions extended to the right. Similarly, negative y positions extended downward and positive y positions upward (see Figure 1B). This coordinate reference plane was used for all subsequent analyses including for velocity, in which velocity bins referred to the speed and direction of the rat’s trajectory. We used negative velocity to indicate that the rat was moving in the negative x direction (to the left of the arena) or y direction (downward), not that the rat was slowing down. Therefore, our measurements of velocity in x and y represented the same information as the combination of speed and head direction canonically studied in prior works (Hardcastle et al., 2017; Sargolini et al., 2006).

### Using Gaussian Processes to increase data sampling in 4D behavioral space

To examine how velocity and position tuning curves might interact, we considered the full 4D space of 2D position and 2D velocity. We constructed 4D tuning curves by binning the data into 2D position and 2D velocity bins. That is, we treated the following as independent behavioral variables: 1) position in x, 2) position in y, 3) velocity in x, and 4) velocity in y. Based on the coverage of each variable within the open field (Figure 1C), we used bin sizes that would be small enough to allow visualization of dynamics across the behavioral space without yielding so many bins as to become too computationally cumbersome. We settled on 5 cm bins for the two position variables and 10 cm/s bins for the two velocity variables.

Examining the rats’ trajectories throughout the arena showed relatively complete coverage of the open field during foraging sessions (Figure 1C). Based on this even sampling, we computed 2D position tuning curves marginalized over velocity. We also examined how much time the rats spent at different velocities ranging from −35 cm/s to +35 cm/s in both x and y velocity components by constructing 2D histograms (Figure 1D). This visualization of time spent in each 10 cm/s-wide velocity bin revealed dominance of time spent at lower speeds. However, since there was a large drop in occupancy from velocity bins centered at 20 cm/s to those centered at 30 cm/s, we eliminated the outer bins of +/-25 to +/-35 cm/s from consideration, yielding 5 bins across each x- and y-velocity.

Binning the data across 4 variables, or dimensions, revealed that despite the relatively even spatial coverage of different x, y positions in the rats’ trajectories, the majority of bins were not or barely visited across all behavioral sessions (Figure S1). Given that many bins were not visited, it was difficult to estimate the tuning curve at each point in this 4D space. Notably, our bin sizes of 5 cm for position and 10 cm/s for velocity yielded 30 × 30 × 5 × 5 = 22,500 bins. Averaging the neural activity at each recorded point in behavioral space left gaps at points where the rat never experienced a particular position vector at a particular velocity vector. We constructed a 2D histogram for each velocity bin showing the amount of time each position point in the arena was visited (Figure 1E), and in three of the behavioral sessions, rats visited fewer than 50% of the bins (with the exception of the one session, Rat R Day 1, which visited > 90% of the bins).

To capture the full dynamics of neural activity across this 4D space, we turned to methods that could statistically enrich the dataset. Gaussian Processes (GPs) are a statistical inference technique that estimates a mean function over a domain where we have partial, but not full, observations. While previously useful for certain machine learning problems, GPs have only recently been explored as a method for improving sampling of sparse data in the field of navigation (Rule et al., 2023; Ye and Wessel, 2024; Mainali et al., 2025). Here, we used GP regression to obtain a mean estimate of the points in position by velocity space where data was sparse due to the difficulty of sampling all points in the 4D behavioral space. In addition to estimating the mean of functions consistent with our observation points, our GP model also provided the associated variance, allowing us to assess the confidence of the model in its estimate at different points in space. We started by using a simple kernel, the Matérn kernel with *ν* = 5/2, which features an exponential decay as two points become more distant from each other. We did a local parameter search for a good initial value and initialized to a length scale of 10 for each variable and variance of 0.3. The gpflow Python package we used to further optimize these parameters to find the best fit for the data.

Using the Matérn kernel, we found that we could reliably (fraction of variance explained [FVE] > 0.5) reconstruct the 2D position tuning curve of a grid cell when the model was given 50% of the points in training (Figure 2A). We evaluated the performance of the model by computing the FVE on held out data. We trained models with increasing proportions of the data used in training and found that using 25%-50% of the data to train the model yielded a comparable FVE to training on 75% of the data. This indicates that our GP model is capable of explaining held out data well, and thus we proceeded with the Matérn kernel for subsequent analyses.

**Figure 2:**
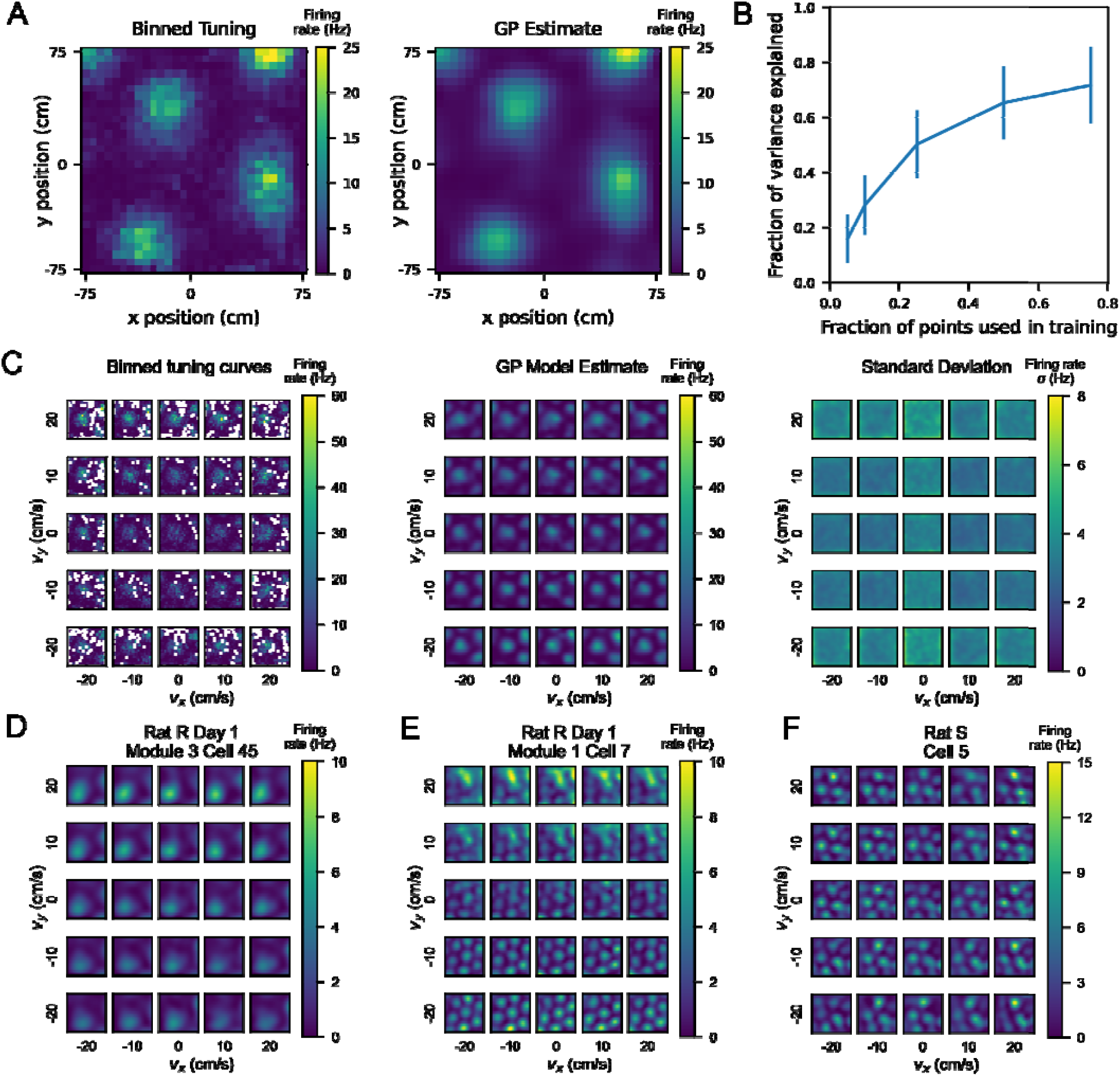
A Gaussian Process (GP) model can estimate firing rates at points in 4D position-velocity space, including those not sampled by the rat’s behavioral trajectory. A) Left: binned firing rate over 2D position for an example neuron, averaged over the entire session, color coded for maximum (yellow) and minimum (blue) values. Bin size is 5 cm. Right: GP estimate, using a simple squared exponential kernel, of the firing rate at the same bin size trained on 50% of the data used to generate the averaged tuning curve on the left. B) Fraction of variance explained (FVE) by GP models trained with varying proportions of the tuning curve.10 different random splits of the data were used to train a model on each cell at different proportions, and the FVE is shown as the mean +/-standard deviation across all 1490 samples (149 cells, 10 iterations each). C) Left: 4D tuning curve for an example neuron, averaged over one behavioral session. As in Figure 1E, each square represents the position tuning curve for the neuron at one particular x- and y-velocity bin. The position tuning curve is represented by a heatmap color coded for maximum (yellow) and minimum (blue) values of firing rate (Hz). Middle: The estimate of the 4D tuning curve given by the GP model for this neuron. Right: The standard deviation given by the GP model as a measure of confidence for each point in the estimate. Heatmap is color coded for maximum (yellow) and minimum (blue) values of standard deviation of firing rate (Hz). D, E, F) Additional examples of single cell tuning curves estimated by the GP model, across different behavioral sessions and rats. Different neurons showed different dependence of position tuning on velocity. Heatmaps are color coded by maximum (yellow) and minimum (blue) values of firing rate in Hz, as in (C).

### Single cell tuning curves show velocity dependence of position tuning

To consider the interaction of position with velocity in neural coding, we examined single cell position-velocity tuning curves. Single cell 4D tuning curves were estimated by the GP model, with each cell modeled independently from all other cells (Figure 2C). Notably, GP regression relies on the proximity of observed points to enhance confidence in estimates. In our case, these points were bins in 4D position-velocity space that had been visited at least 10 times (Figure S1A and B), and we estimated the interpolation of missing data for all other bins. Here, GP works well in part because all cells were recorded simultaneously for a given session, meaning that the behavioral statistics are identical for all of the recorded cells in that session. Moreover, prior work has shown that the grid cell population can be classified into modules, in which all grid cells in a given module share the same orientation (Hafting et al., 2005) and grid spacing (Stensola et al., 2012). Our GP model is instantiated on each cell independently, allowing for cell-to-cell comparisons across model performance on similar behavioral statistics.

As seen in Figure 2B, from the binned 4D tuning curves, we trained the GP model to estimate the firing rate at all points in 4D space, including at points not sampled by the rat. The standard deviation estimate remained fairly low across the middle of the arena in most velocity bins but was higher in the edges and corners, where the rat was least likely to explore at high speeds (Figure S1C).

We next examined if position tuning varied from one velocity bin to another. One possibility is that grid cell tuning may differ between low and high speeds and/or different directions of movement. This would indicate that the animal’s internal map of space can change depending on the direction and speed of movement. Although prior work has observed that grid cells increase their signal-to-noise ratio (Hardcastle et al., 2017) with increases in speed, our approach allowed us to disentangle an increase with firing rate across unchanging grid fields from a change in the properties of the fields themselves as a result of velocity.

We first considered all grid cells within a given module for a given recording session. For each cell, we calculated the cosine similarity of the 2D position tuning curve marginalized over all velocity with position tuning curves restricted to specific velocity bins. We then averaged these cosine similarities across cells in each module. Using this similarity metric, we found that as velocity increased to more extreme values, grid cell position tuning curves became less similar to their tuning marginalized over all velocities (Figure S2A). Because cosine similarity is agnostic to differences in the magnitude of firing rates, this distribution of values indicates that position tuning is not uniform across velocity bins.

Next, we considered how individual grid cell position tuning curves might change across different velocity bins (Figure 2D-F). We found that some cells exhibit more gradual changes in position tuning across velocity bins with grid fields largely in the same positions (Figure 2B center, Figure S2A). Other cells showed different patterns across velocity bins (Figure S2B), including differences in the edges of the grid fields (2D), gain changes modulated by velocity (2E), and deterioration of the grid fields in part of the arena (2F). We found that these differences in velocity dependence exist across cells even within the same module (Figure S3A), and other cells in these modules did not exhibit velocity-dependent tuning to the same degree (Figure S3B).

We confirmed this finding was not an artifact of our GP model by simulating a spike train using a null model with invariant tuning drawn from the zero-velocity bin of each cell (Figure S4). Our null model showed that the non-separability is observable from our GP analysis only when there is enough coverage of the behavioral space, as in Rat R Day 1, unlike in the other behavioral sessions. This is consistent with our finding that the GP model does not outperform the separable model on the other three behavioral sessions.

We also examined whether grid field parameters changed systematically across speed alone, as this could explain the velocity-dependent changes we saw. For this analysis, we considered data from Rat R Day 1, as our previous analysis demonstrated this session as the most robust in terms of coverage of the behavioral space. However, the distributions of grid score and grid scale did not differ across speed bins in Rat R, Day 1 (Figure S5A-B). Even so, we did see some clustering in grid scores across velocity space on an individual cell basis (Figure S5C).

### Using a separable model to examine velocity gain modulation on position tuning

Given the velocity dependence of position tuning we saw in many cells’ tuning curves, we wondered if this dependence could reflect a gain modulation on position tuning by velocity, a coding feature previously observed in 2D tuning curves (Hardcastle et al., 2017). Mathematically, this would be equivalent to the firing rate as a function of 4D position-velocity being factorizable into two functions: one purely a function of 2D velocity and another purely a function of 2D position. To consider whether gain modulation by velocity influenced position tuning, we examined whether a model trained on this assumption of factorizability could capture most or all of the variance explained by our full 4D model. We used gradient descent to optimize a separable model as a product of two functions (Figure 3A), one over position space and another over velocity space. We then compared the fraction of variance explained (FVE) by this separable model on 20% held-out data to that of our GP model.

**Figure 3:**
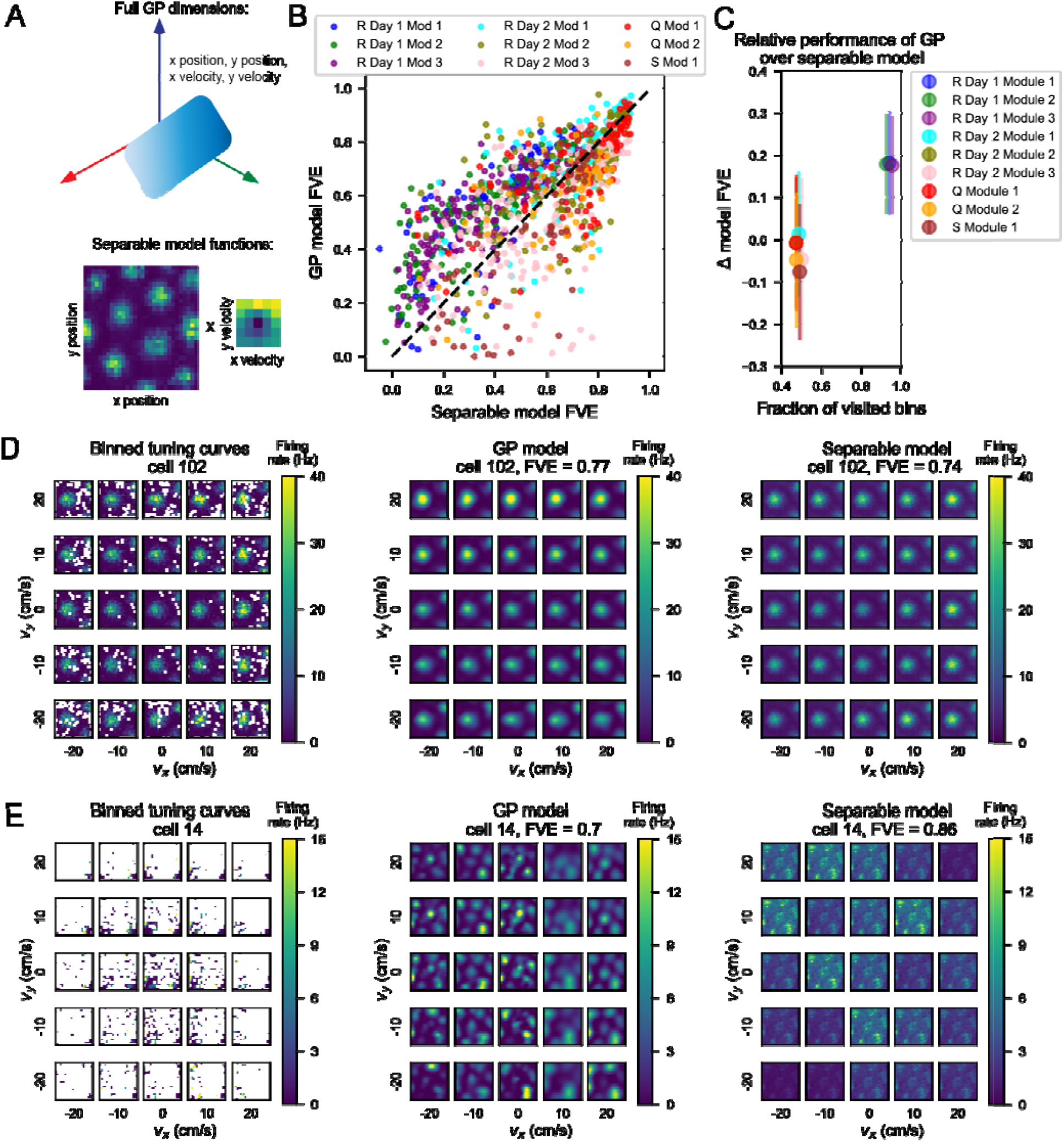
Relative performance of a separable model to the full GP model depends on the amount of data present in a given behavioral session. A) Schematic describing how the separable model differs from the full GP model. Top: Illustration of the GP model, where the blue gradient represents a 4D function over 2D position and 2D velocity. Bottom: Illustration of the two functions optimized by the separable model, one over position space (left) and one over velocity space (right). B) Scatterplot of performance between the separable GP model and the full GP model. We computed fraction of variance explained (FVE) on held-out data as a measure of performance for each model. In Rat R Day 1, points lay generally above and to the left of the unity line, showing that the full model performs better than the separable model. Each point represents a neuron, color coded by rat (R, Q, or S), session (day) and module (number). C) Comparison of the occupancy of behavioral space across one session to performance of the separable model. Data were jittered so all points could be observed. Circles represent mean, and lines indicate standard deviation. Session-to-session variability depended on amount of data collected. The x-axis shows bins in position-velocity space where the rat visited at least once. For each behavioral session, the fraction of visited bins out of all bins is plotted against the average FVE (on held out data) of the separable model. D) The GP model and separable model both performed well on cells with separable tuning. An example cell (102) from Rat R, Day 1, Module 3 in the upper right region of the session (purple) point cloud (panel B), indicating relatively good performance from both the separable and full GP models. Left: binned 4D tuning curve averaged over recorded data. Center: estimate of the 4D tuning curve by the GP model with a fraction of variance explained of 0.77. Right: estimate of the 4D tuning curve by the separable model with a fraction of variance explained of 0.74. E) In sessions with less data, the separable model may generalize better than the full GP model. An example cell from Rat Q, Module 1, on the right side of the point cloud, indicating better performance from the separable model than from the full GP model. Left: binned 4D tuning curve averaged over recorded data. Center: estimate of the 4D tuning curve by the full model with a fraction of variance explained of 0.7. Right: estimate of the 4D tuning curve by the separable model with a fraction of variance explained of 0.86.

To do this, we plotted FVE of the separable model on the x-axis against FVE of the full model on the y-axis and found that there was variability from session to session in terms of where the point cloud lay (Figure 3B). In cases where a point lay to the left and above the dashed unity line, the GP model captured more variance than the separable model. The leftover variance not captured by the separable model must therefore be an interaction between the position and velocity variables, and the representation of 2D position and 2D velocity in grid cells is necessarily 4D. In modules recorded in Rat R on Day 1, points lay mostly to the upper left of the unity line, but other sessions showed more spread of points across the unity line as well. We noticed that the behavioral sessions varied by data density, and a higher proportion of bins visited by the rat correlated with a higher Δ FVE as GP model FVE - separable model FVE (Figure 3C).

We also observed a range of FVE for both models within behavioral sessions and modules. We investigated the tuning curves of cells in the upper right corner of our scatterplot (Fig. 3B). Using cell 102 from rat R module 2 day 1 as an example, we plotted the binned 4D tuning curves, the estimate derived from the GP model (FVE = 0.77), and the estimate from the separable model (FVE = 0.74) (Figure 3D). Both tuning curves contained grid fields in roughly the same locations as in the binned tuning curves, with an average cosine similarity of 0.94 across models and SEM of 0.002.

We also examined cells where the separable model outperformed the GP model (Figure 3E). Using cell 14 from Rat Q Module 2, we again plotted the binned tuning curves next to the estimates from the GP models. The separable model captured the neural data with FVE of 0.86, whereas the full model performed a little worse with FVE of 0.7. One possibility is that the data sparsity shown in the left panel of Figure 3E contributed to the relative performance of the separable model, especially as we could only evaluate performance on bins visited by the rat. We noticed that in Rat R Day 2, Rat Q, and Rat S sessions, which were each much shorter in session length than Rat R Day 1 (38, 52, and 36 as opposed to 131 minutes), there was no significant difference in model performance or the separable model performed slightly better, while in most cases the GP model outperformed the separable model in Rat R Day 1 (Figure 3C, S3A). Because the data observed were clustered in position-velocity space (Figure 3E left), the separable model may perform better than the GP model because it has fewer parameters to tune. In addition to this large split in model performance across sessions, there was variable model performance within a given session across individual cells.

We considered what factors might contribute to this spread in performance within sessions. We noticed that mean firing rate was correlated with performance of the GP model (all *p* < 0.01), as well as with performance of the separable model (all *p* < 0.01, Figure S6B). Although we were able to find the best fit line for each session modeled using GPs, we observed that some of the sessions modeled by the separable model saturated at FVE of 1 at a lower mean, and these tended to be sessions with little data (Figure S6B right). We next asked if there was a significant trend of Δ FVE (GP model FVE - separable model FVE) with mean firing rate and found a significant negative correlation (*p* < 0.01) in all sessions except Rat S, where the correlation was not significant (*p* = 0.08). This negative correlation is driven by the early saturation of mean firing rate on separable model FVE, particularly in sessions with less data.

Given that one model separates position and velocity tuning and the other does not, we wanted to characterize whether the GP model was truly capturing non-separability of tuning in cells. To do so, we examined the heatmaps of cosine similarity of various velocity bins with the 2D position tuning curve. Non-separability of tuning curves could be characterized by grid fields translating in location across velocity bins or individual fields disappearing or appearing independently of other fields. Scaling of firing rate magnitude of position tuning curves across velocity bins would yield a separable tuning curve that would also result in a similar cosine similarity value across bins. Thus, we considered the standard deviation in cosine similarity (SDCS) across velocity bins to evaluate how non-separable a given cell’s velocity and position tuning is. We compared Δ FVE (GP model FVE – separable model FVE) to the corresponding SDCS across cells (Figure 4B), finding a significant positive correlation. We created a Linear Mixed Effects Model (LMEM) to understand the contribution of various factors to Δ FVE. We found that several of these factors significantly influenced Δ FVE, including module (*β* = −0.002, p = 2e-5), mean firing rate (*β* = −0.044, p = 2e-62), SDCS (*β* = 4.76, p = 4e-75), and data density as proportion of bins visited (*β* = 0.573, p = 8e-218). Given these statistics, we examined individual cell tuning curves and found that there were large (>0.2) differences in FVE when grid fields appear and disappear across different velocity bins, which corresponds to a higher SDCS (Figure 4C-H).

**Figure 4:**
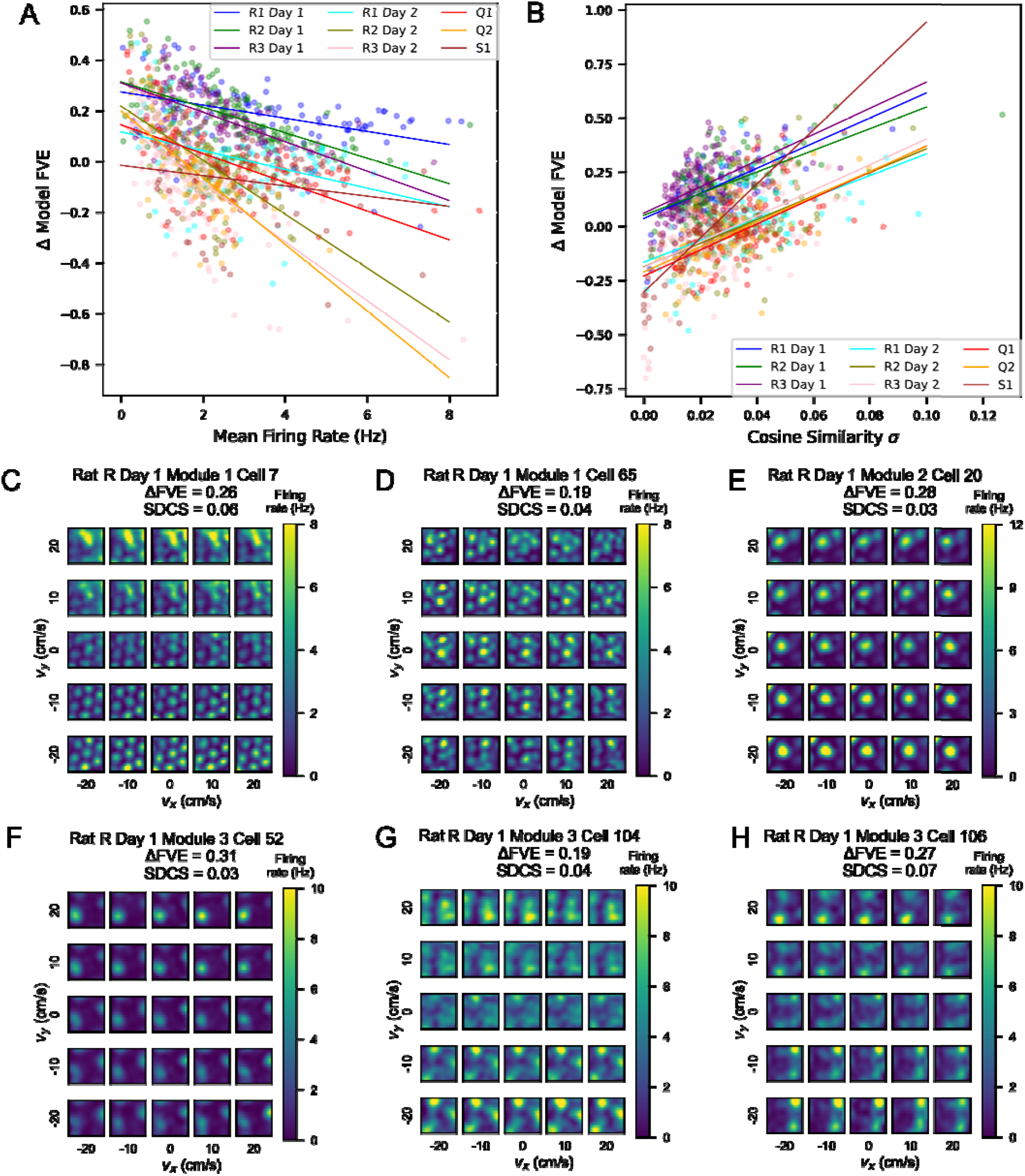
Many cells show inseparable tuning between position and velocity, leading to better performance by the full GP model. A) Scatterplot showing the difference in FVE between the full GP model and the separable model against mean firing rate in Hz. Each point represents a cell, color-coded by rat, module and session. The best fit linear regression lines are plotted in the same colors. B) Scatterplot showing the difference in FVE between the full GP model and the separable model against the standard deviation in cosine similarity (SDCS) of position tuning curves in each velocity bin with the marginalized 2D position tuning curve over all velocities. As in (A), each point is a cell, and the best fit regression lines are plotted in the same colors. C-H) Examples of inseparable tuning curves from Modules 1, 2, and 3 of Rat R Day 1. In each panel, the estimate of the GP model is visualized, with annotations for the FVE of the GP model and the standard deviation of the cosine similarity (SDCS) as a measure of non-separability for each cell.

We also asked whether the grid score or grid scale of a given cell was correlated with the difference in performance of the GP and separable models. We did not find a significant correlation between these variables and Δ FVE (Figure S3B-C). Thus, we concluded that the GP model outperforms the separable model when there is enough data to observe non-separable tuning curves and is not dependent on the grid scale or grid score.

## Discussion

Grid cells in MEC have traditionally been examined in terms of their position tuning curves in 2D environments, which depict the firing rate of a given grid cell across the x and y axes of space (Fyhn et al., 2004; Hafting et al., 2005). Similarly, speed tuning has traditionally been examined using 1D tuning curves that depict average firing rates as a function of an animal’s running speed (Hinman et al., 2016; Kropff et al., 2015). While these approaches have yielded foundational insight into grid and speed cell coding properties, they have not fully addressed whether grid cells encode position and velocity independently or whether grid cells integrate these variables in a non-separable manner. To address this question, we considered the tuning curves in 4D behavioral space, across both 2D position and 2D velocity. Leveraging an existing dataset from Gardner et al., 2022, which includes long behavioral sessions of random foraging and clearly defined grid modules, we analyzed how 2D position tuning curves shifted across different velocities.

Given the challenge of sampling all possible combinations of velocities, we binned our velocity data into 10 cm/s bins and averaged the firing rate within each velocity bin. The 10 cm/s width of our velocity bins allowed us to avoid bins with zero observations in the 2D velocity tuning curve. Still, the 4D space was sparsely sampled, particularly in high-speed or edge-of-arena conditions. To overcome this limitation, we used GPs to estimate information at unsampled points in 4D behavioral space. This approach built upon previous work utilizing GPs to model grid cell tuning curves (Rule et al., 2023; Ye and Wessel, 2024). While GPs have been used to make inferences in 2D position space (Rule et al., 2023) or understand the neural manifold of the population at different speeds (Ye and Wessel, 2024), our approach extended their use to full 4D position-velocity space. By applying GPs to estimate position-velocity tuning curves in the 4D space on a cell-by-cell basis, we were able to uncover non-separability between the tuning curves for position and velocity. Our finding of non-separability in single cell 4D tuning curves indicates that position and velocity coding may change jointly, rather than independently. In other words, position tuning in MEC may flexibly shift depending on velocity, suggesting a higher-dimensional representation of space that could support navigation across different velocity states.

Drawing from studies investigating separable space-time coding in sensory neuroscience (Cowan et al. 2016; DeAngelis et al. 1993; Mazer et al. 2002; Depireux et al. 2001), our definition of separability was specifically narrow. In essence, it is equivalent to understanding whether these 4D tuning curves could be factorized into a velocity term and a position term, to understand whether neurons could compute velocity and position tuning independently. Moreover, this 4D tuning points to the importance of considering grid cell activity as a joint function of multiple behavioral variables, rather than considering each behavioral variable completely independently. It will be interesting for future work to examine the possible mechanisms underlying velocity-associated changes in position coding, such as the appearance or disappearance of grid fields.

Additional work to elucidate the causes of the non-separability we observed would further improve our understanding of multi-dimensional coding in MEC. There are many possible reasons for non-separability in position-velocity coding, including the presence of prospective or retrospective coding in this part of the navigational circuit (Campbell et al. 2021; Kropff et al. 2015). In this way, the directionality and speed of motion could impact the representation of space in grid cells. We speculate that such mechanisms might arise from traveling waves within the circuit. Other sources of non-separability include left-right theta sweeps modulated by speed (Vollan et al. 2025). Our work here shows that there are differences in grid cell position tuning based on velocity and does not exclude these mechanisms, but further work is needed to understand the origin of this non-separability.

Previous work has shown that grid cells in the MEC are modulated by changes in task demands or reward, but it is not clear how much of this modulation reflects changes in behavioral statistics between tasks or reward contexts (Butler et al., 2019; Boccara et al., 2019). For example, a recent theoretical study found that shifts in grid cell coding could result from changes in an animal’s behavioral statistics (Nayebi et al., 2021). Our findings complement this idea by raising the possibility that when behavior changes, grid cells change their coding properties due to the intrinsic velocity dependence of the grid cell position code. Future work could aim to further investigate this by precisely manipulating behavioral variables, like velocity, and task variables, such as random versus goal-directed foraging, while modeling the high-dimensional influence of these variables on neural activity.

Our examination of the conjunctive tuning of MEC grid cells across position and velocity variables represents an initial foray into the complex field of examining high-dimensional spatial representations. However, this framework can be easily extended to include additional behavioral axes. For example, future studies could incorporate variables such as pupil size or whisking, along with position and velocity. This work illustrates one way in which such high-dimensional representations can help identify interaction factors that would otherwise be averaged out.

## Acknowledgements

We thank Charlotte Herber, Ben Sorscher, Dan Biderman, and Scott Linderman for helpful discussions in early stages of the project; Charlotte Herber, Emily Aery Jones, and Alex Gonzalez for sharing code; and Tom Clandinin and Shaul Druckmann for feedback throughout the project. LMG is an HHMI Investigator and this work was supported by an 1R01MH126904-01A1 (LMG), R01MH130452 (LMG), BRAIN Initiative U19NS118284 (LMG), P50 DA042012 (LMG), The Vallee Foundation (LMG), The James S. McDonnell Foundation (SG, LMG), The Simons Foundation 542987SPI (SG, LMG) and the NSF CAREER and Schmidt Foundation (SG), Stanford Graduate Fellowship (LJW), and the NSF Graduate Research Fellowship (LJW).

## Author Contributions

Conceptualization (LJW, SG, LMG), Methodology (LJW, SG, LMG), Investigation (LJW), Visualization (LJW), Funding acquisition (SG, LMG), Project administration (SG, LMG), Supervision (SG, LMG), Writing – original draft (LJW, LMG), Writing – review and editing (LJW, LMG).

## Data and Code Availability

Data is available as provided by Gardner et al. 2022 at https://figshare.com/articles/dataset/Toroidal topology of population activity in grid cells/16764508. Code is available upon reasonable request.

## Methods

### Data Processing

We used behavioral and neural data from Gardner et al. 2022, extracted using the first steps of their utility code. This provided x- and y-coordinates of position, as well as spike count over the time each rat spent exploring the open arena. All neural data corresponded to identified grid cells and had been clustered into modules based on grid spacing. One rat (R) was recorded over multiple days and featured three co-recorded modules, and another rat (Q) featured two co-recorded modules. Only one identified grid module was recorded from rat S. On average, there were more than 100 grid cells per module and day. We modeled each cell independently.

We computed velocity by finding the difference between adjacent time points in the smoothed position trajectory for both x and y. This allowed us to construct a vector of x-position, y-position, x-velocity, and y-velocity over time. The spikes were sorted into 10-ms-wide time bins. We did not consider points where the x or y velocity of the rat was greater than 50 cm/s.

### Binning

We computed position tuning curves empirically by binning the environment in the x and y directions into 5 cm bins for a total of 30 bins. We then found the mean firing rate of each neuron within a particular bin over the course of the entire session by dividing the sum of spikes by time spent in that position bin. We extended this to 4 dimensions by splitting each position bin into further x- and y-velocity bins. Due to occupancy constraints, we limited our analysis to velocities between −25 and +25 cm/s in both x and y directions and then binned these velocities into 5 bins of 10 cm/s each. Thus, our 4D tuning curve had dimensions 30 by 30 by 5 by 5, for each neuron, given by:

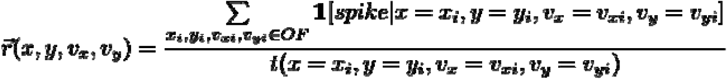

where *t* reflects the occupancy of the animal in each bin. These binning procedures allowed us to compute our 2D histograms of occupancy across 2D velocity and 2D position space, as well as our arrays of 2D histograms representing occupancy in full 4D space.

### Gaussian Processes Regression

We used Gaussian Processes (GP) regression to estimate tuning curves across 4D behavioral space where occupancy was low due to behavioral constraints. In order to estimate the 4D tuning curve of average firing rates across bins, we used a GP given by:

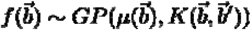

Here we use *b* to denote the 4D behavior vector of x-position, y-position, x-velocity, and y-velocity at a particular point. We used the gpflow package in Python to estimate the full 4D tuning curve as well as the standard deviation of this estimate. We computed a different model and estimate for each cell. We used GPUs to accelerate this computation. The outputs of this model were two 4D matrices of the same size as the binned tuning curves as an estimation of a function over the range of position and velocity bins.

We tested a variety of kernels *K*(*b, b*^⍰^) for comparison, including Matérn kernels with nu = 1/2, 3/2, and 5/2, as well as the Radial Basis Function (RBF). Because the results were similar across different values of the nu parameter, we chose nu = 5/2 to be the standard value across our results. We initialized the length scale parameter to 10 cm for the x and y position dimensions and 10 cm/s for the x and y velocity dimensions, but the gpflow package we used to train our model optimized these lengthscale parameters (with a scipy optimizer) to find the best fit. For each cell, we used a k-fold cross-validation procedure to produce k sets of training and validation data. For each of the k sets, we initialized a GP model with the above kernel parameters as well as with training data. Then we optimized the GP model using scipy.optimizer, which minimized the negative log likelihood. In order to assess the performance of each model, we computed the Fraction of Variance Explained (FVE) on the held-out validation data:

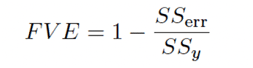

where *SS*_err_ is the variance of the residuals and *SS*_y_ is the variance of the held-out data. Lastly, we computed the estimate and variance of the GP model prediction at each point in 4D space. In subsequent analyses, we averaged the FVE and outputs of the model over the k folds of our analysis.

### k-fold Cross-Validation Procedure

We wanted to provide training points for the GP models at points in behavioral space that had been visited semi-frequently. Based on the density of data at various occupancy thresholds (Figure S1A), we set a threshold of 10 measurements within a given position-velocity bin to be considered “high-occupancy.”

Among these high-occupancy bins, we set k = 5 and computed an 80/20 training/validation split on the stable points. We then averaged over the estimate for each model to produce a cross-validated inference matrix over each cell. We computed the fraction of variance explained for each estimate on the held-out validation set and averaged these scores as well for each neuron. When comparing fraction of variance explained across models, we used the same sets of points as training, validation, and test points.

### Separable model optimization

We constructed our separable model as the outer product of two matrices, one for position space and the other for velocity space. Thus, our model prediction was determined by

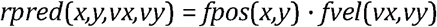

We used gradient descent implemented through PyTorch to minimize the mean squared error between the predicted firing rate and observed firing rate as a loss function:

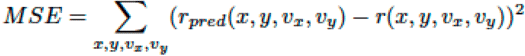

We also implemented a non-negativity constraint on the matrices using projected gradient descent, such that the gradient descent step was taken and then the matrices were projected back to the nonnegative domain.

### Non-separability analysis

To examine the non-separability of a given cell’s 4D tuning curve, we found the cosine similarity between the 2D position tuning curve at each velocity bin *r*(*x,y,v*_x_,*v*_y_) and the marginalized 2D position tuning curve, 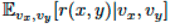.

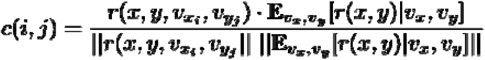

This yields a 5×5 matrix of cosine similarity scores to the marginalized 2D position tuning curve. We found the standard deviation of the cosine similarity values (SDCS) as a measure of the non-separability of single cell tuning curves.

### Analysis of Grid Cell Properties

We followed the standard procedure for calculating grid score and grid scale. We computed the autocorrelation of the rate map and found the locations of the six closest firing field peaks relative to the center peak. We determined the grid scale by finding the average distance from each of the six peaks to the center peak, and we determined the grid score by finding the difference between the mean correlation with a rotated map at 60^⍰^ and 120^⍰^ compared to 30, 90, and 150^⍰^.

### Linear regression of cell properties with model performance

We asked whether these cell properties, including mean firing rate, grid score, standard deviation of cosine similarity, and grid scale, correlated with model performance. Using the linregress method from scipy.stats package, we computed the best fit line relating a vector of each property across cells to the FVE of a given model or the difference in the FVE, Δ*FVE*.

### Linear Mixed Effects Models

We leveraged linear mixed effects models (LMEMs) to understand the contribution of various factors to the difference in FVE Δ*FVE* = *FVE*_*GP*_ - *FVE*_separable_. We implemented these models with the mixedlm method of the statsmodels.api package and treated animal identity as a random effect. We incorporated continuous variables of module, mean firing rate, data density, and standard deviation of cosine similarity (SDCS) as fixed effects. We computed data density as the fraction of bins in 4D space visited by the animal and SDCS as described above. Our LMEM converged using restricted maximum likelihood estimation.

### Validation of GP Model with Simulated Data

In order to ensure that our GP model was not hallucinating firing fields, we constructed a null model by isolating the 2D position tuning curve *r*_*0*_ at the zero-velocity bin (centered at v_x_ = 0 and v_y_ = 0) and generating a spike train along the observed behavioral trajectories using a Poisson model. We then ran our GP model on this simulated data, generating the full 4D tuning curve. Given that these data were generated conditionally on *r*_*0*_, the 4D tuning curves produced by the GP model should not produce position tuning curves that differ across velocity bins. We computed the SDCS score on these null model 4D tuning curves and compared this score to the corresponding SDCS score computed on the actual data in order to determine whether the model was incorrectly generating non-separability when there was none.

## Supplemental Materials

Supplemental Figure 1: Behavioral statistics by session illustrate the need for data enrichment.

Supplemental Figure 2: Standard deviation of cosine similarity (SDCS) shows velocity dependence of position tuning and differs across cells.

Supplemental Figure 3: Additional examples of individual cells with non-separability in Rat R, Day 1.

Supplemental Figure 4: Analysis of GP performance on simulated data.

Supplemental Figure 5: Additional characterization of non-separability on neurons within the top 50% of SDCS per module.

Supplemental Figure 6: Differences in the performance of the GP and separable models are driven by mean firing rate but not grid score or grid scale.

## References

Fyhn, M., Molden, S., Witter, M. P., Moser, E. I., & Moser, M.-B. (2004). Spatial Representation in the Entorhinal Cortex [Publisher: American Association for the Advancement of Science]. Science, 305(5688), 1258–1264. 10.1126/science.1099901

Hafting, T., Fyhn, M., Molden, S., Moser, M.-B., & Moser, E. I. (2005). Microstructure of a spatial map in the entorhinal cortex [Publisher: Nature Publishing Group]. Nature, 436(7052), 801–806. 10.1038/nature03721

McNaughton, B. L., Battaglia, F. P., Jensen, O., Moser, E. I., & Moser, M.-B. (2006). Path integration and the neural basis of the ‘cognitive map’ [Publisher: Nature Publishing Group]. Nature Reviews Neuroscience, 7(8), 663–678. 10.1038/nrn1932

Sargolini, F., Fyhn, M., Hafting, T., McNaughton, B. L., Witter, M. P., Moser, M.-B., & Moser, E. I. (2006). Conjunctive representation of position, direction, and velocity in entorhinal cortex. Science (New York, N.Y.), 312(5774), 758–762. 10.1126/science.1125572

Guanella, A., Kiper, D., & Verschure, P. (2007). A model of grid cells based on a twisted torus topology. International Journal of Neural Systems, 17(4), 231–240. 10.1142/S0129065707001093

Burak, Y., & Fiete, I. R. (2009). Accurate Path Integration in Continuous Attractor Network Models of Grid Cells [Publisher: Public Library of Science]. PLOS Computational Biology, 5(2), e1000291. 10.1371/journal.pcbi.1000291

Stensola, H., Stensola, T., Solstad, T., Frøland, K., Moser, M.-B., & Moser, E. I. (2012). The entorhinal grid map is discretized. Nature, 492(7427), 72–78. 10.1038/nature11649

Kropff, E., Carmichael, J. E., Moser, M.-B., & Moser, E. I. (2015). Speed cells in the medial entorhinal cortex. Nature, 523(7561), 419–424. 10.1038/nature14622

Hinman, J. R., Brandon, M. P., Climer, J. R., Chapman, G. W., & Hasselmo, M. E. (2016). Multiple Running Speed Signals in Medial Entorhinal Cortex. Neuron, 91(3), 666–679. 10.1016/j.neuron.2016.06.027

Hardcastle, K., Maheswaranathan, N., Ganguli, S., & Giocomo, L. M. (2017). A Multiplexed, Heterogeneous, and Adaptive Code for Navigation in Medial Entorhinal Cortex. Neuron, 94(2), 375–387.e7. 10.1016/j.neuron.2017.03.025

Boccara, C. N., Nardin, M., Stella, F., O’Neill, J., & Csicsvari, J. (2019). The entorhinal cognitive map is attracted to goals. Science (New York, N.Y.), 363(6434), 1443–1447. 10.1126/science.aav4837

Spalla, D., Treves, A., & Boccara, C. N. (2022). Angular and linear speed cells in the parahippocampal circuits. Nature Communications, 13: 1907.

Butler, W. N., Hardcastle, K., & Giocomo, L. M. (2019). Remembered reward locations restructure entorhinal spatial maps. Science (New York, N.Y.), 363(6434), 1447–1452. 10.1126/science.aav5297

Low, I. I. C., Williams, A. H., Campbell, M. G., Linderman, S. W., & Giocomo, L. M. (2021). Dynamic and reversible remapping of network representations in an unchanging environment. Neuron, 109(18), 2967–2980.e11. 10.1016/j.neuron.2021.07.005

Nayebi, A., Attinger, A., Campbell, M., Hardcastle, K., Low, I., Mallory, C. S., Mel, G., Sorscher, B., Williams, A. H., Ganguli, S., Giocomo, L., & Yamins, D. (2021). Explaining heterogeneity in medial entorhinal cortex with task-driven neural networks. Advances in Neural Information Processing Systems, 34, 12167–12179. Retrieved April 1, 2025, from https://proceedings.neurips.cc/paper/2021/hash/656f0dbf9392657eed7feefc486781fb-Abstract.html

Gardner, R. J., Hermansen, E., Pachitariu, M., Burak, Y., Baas, N. A., Dunn, B. A., Moser, M.-B., & Moser, E. I. (2022). Toroidal topology of population activity in grid cells [Publisher: Nature Publishing Group]. Nature, 602(7895), 123–128. 10.1038/s41586-021-04268-7

Rule, M. E., Chaudhuri-Vayalambrone, P., Krstulovic, M., Bauza, M., Krupic, J., & O’Leary, T. (2023). Variational log-Gaussian process methods for grid cells [eprint: https://onlinelibrary.wiley.com/doi/pdf/10.1002/hip Hippocampus, 3 1235–1251. 10.1002/hipo.23577

Ye, Z., & Wessel, R. (2024, September). Speed modulations in grid cell information geometry [Pages: 2024.09.18.613797 Section: New Results]. 10.1101/2024.09.18.613797

Mainali, N., Azeredo Da Silveira, R., & Burak, Y. (2025). Universal statistics of hippocampal place fields across species and dimensionalities. Neuron, S0896627325000431. 10.1016/j.neuron.2025.01.017

Cowan, C. S., Sabharwal J., & Wu S. M. Space-time codependence of retinal ganglion cells can be explained by novel and separable components of their receptive fields. Physiol Rep. 2016 Sep;4(17):e12952. doi: 10.14814/phy2.12952. PMID: 27604400; PMCID: PMC5027358.

DeAngelis, G. C., Ohzawa, I., & Freeman, R. D. Spatiotemporal organization of simple-cell receptive fields in the cat’s striate cortex. I. General characteristics and postnatal development. J Neurophysiol. 1993 69:4, 1091–1117

Mazer, J. A., Vinje, W. E., McDermott, J., Schiller, P. H., & Gallant, J. L. Spatial frequency and orientation tuning dynamics in area V1. Proc Natl Acad Sci U S A. 2002 Feb 5;99(3):1645–50. doi: 10.1073/pnas.022638499. Epub 2002 Jan 29. PMID: 11818532; PMCID: PMC122244.

Depireux, D. A., Simon, J. Z., Klein, D. J., & Shamma, S. A. Spectro-temporal response field characterization with dynamic ripples in ferret primary auditory cortex. J Neurophysiol. 2001 Mar;85(3):1220–34. doi: 10.1152/jn.2001.85.3.1220. PMID: 11247991.

Campbell, M. G., Attinger, A., Ocko, S. A., Ganguli, S., & Giocomo, L. M. Distance-tuned neurons drive specialized path integration calculations in medial entorhinal cortex. Cell Rep. 2021 Sep 7;36(10):109669. doi: 10.1016/j.celrep.2021.109669. PMID: 34496249; PMCID: PMC8437084.

Vollan, A. Z., Gardner, R. J., Moser, M. B., & Moser, E. I. Left–right-alternating theta sweeps in entorhinal– hippocampal maps of space. Nature 639, 995–1005 (2025). doi: 10.1038/s41586-024-08527-1

