## Supplemental Figures for "Gaussian Process Inference Reveals Non-separability of Position and Velocity Tuning in Grid Cells"

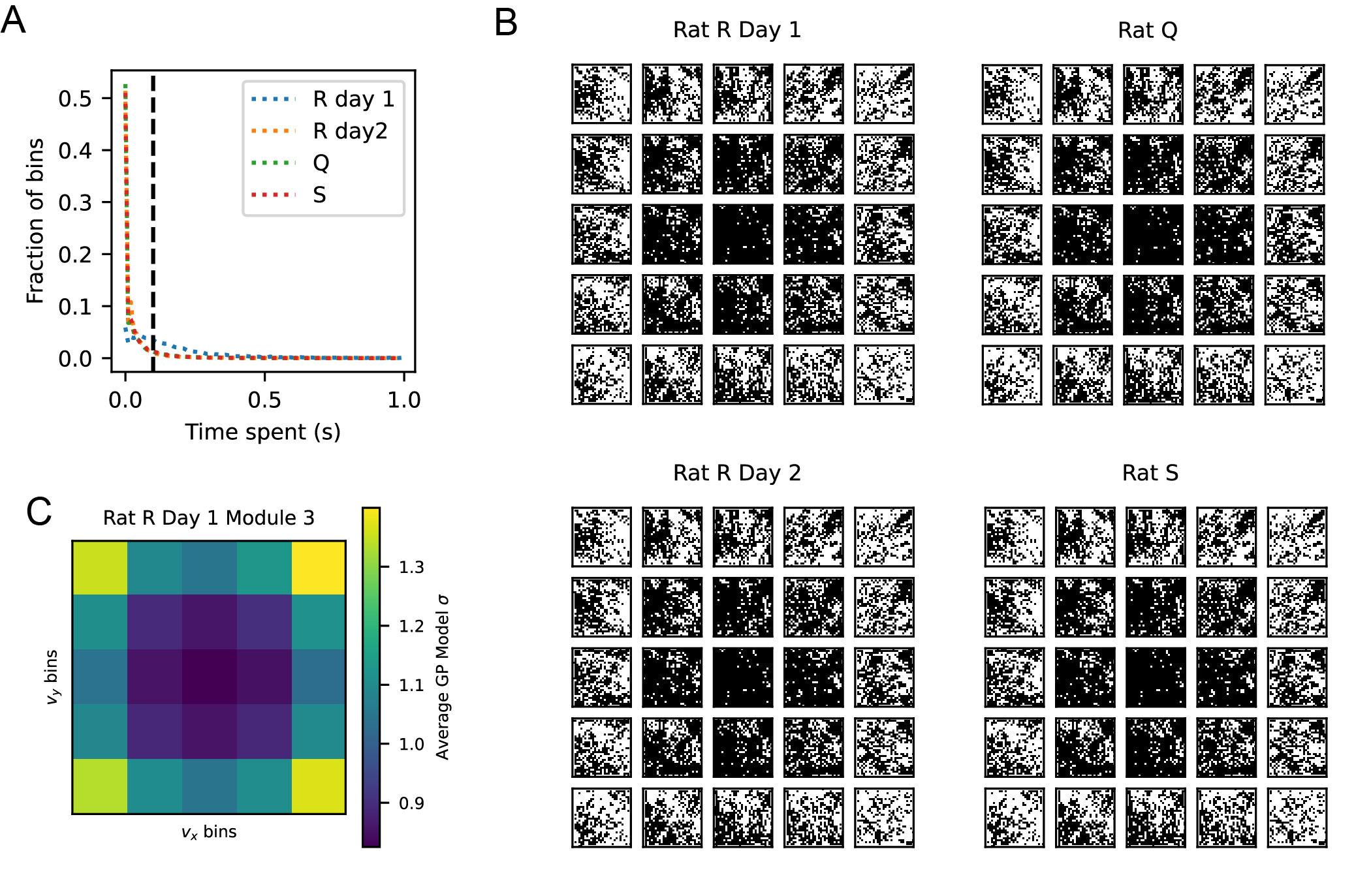

**Supplemental Figure 1: Behavioral statistics by session illustrate the need for data enrichment.**

A) Data density of bins in 4D space for each behavioral session (Rat R on both days, Rat Q, and Rat S). The x-axis depicts the time spent during the behavioral session in a given bin, and the y-axis reports the fraction of bins for a given amount of time spent. Dashed black line indicates our threshold for a “high-occupancy” point, which is at 0.1 s, or 10 time points at 10 ms precision.

B) For each behavioral session, heatmaps with black or white at each bin in 4D space show where the high-occupancy points reside. Black indicates that the bin was visited for at least 0.1 second, and white indicates that it was visited not at all or for less than 0.1 second. C) Mean of standard deviation for the GP estimate in different velocity bins, averaged across position bins and across cells in one session. The standard deviation represents the uncertainty of the GP in its estimate, and it rises toward the edges and corners.

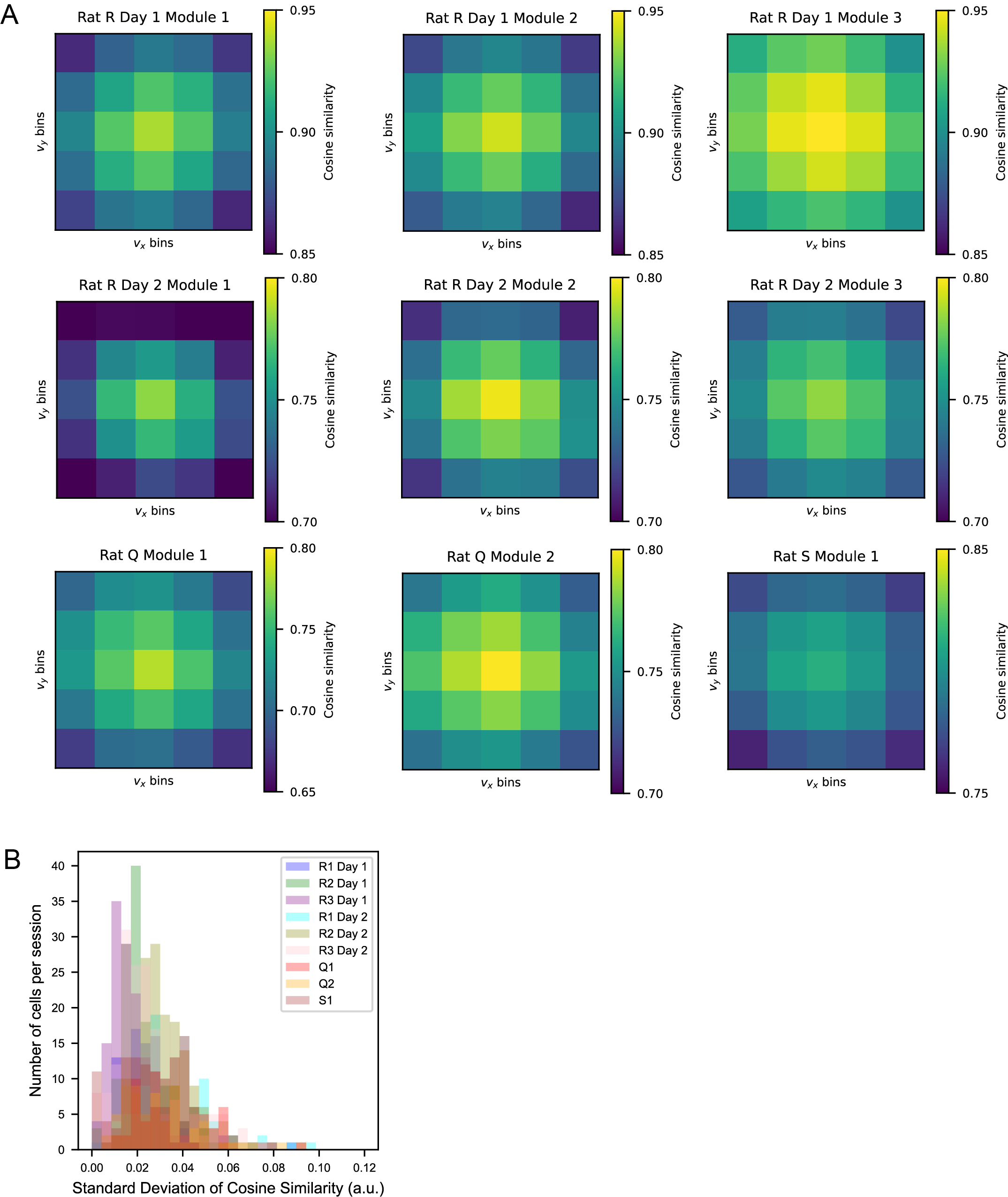

**Supplemental Figure 2: Standard deviation of cosine similarity (SDCS) shows velocity dependence of position tuning and differs across cells.**

A) Cosine similarity of position tuning curve marginalized over all velocities with the position tuning curve at each velocity bin. Each heatmap shows one session. The cosine similarity of each velocity specific position tuning curve to the marginalized position tuning curve is highest in the center bin and decreases to the edges, indicating that there are changes in position tuning as a function of velocity.

B) For each session, histograms of the standard deviation of cosine similarity (SDCS) of the position tuning curve in each velocity bin to the marginalized position tuning curve. A higher value of SDCS indicates more velocity dependence.

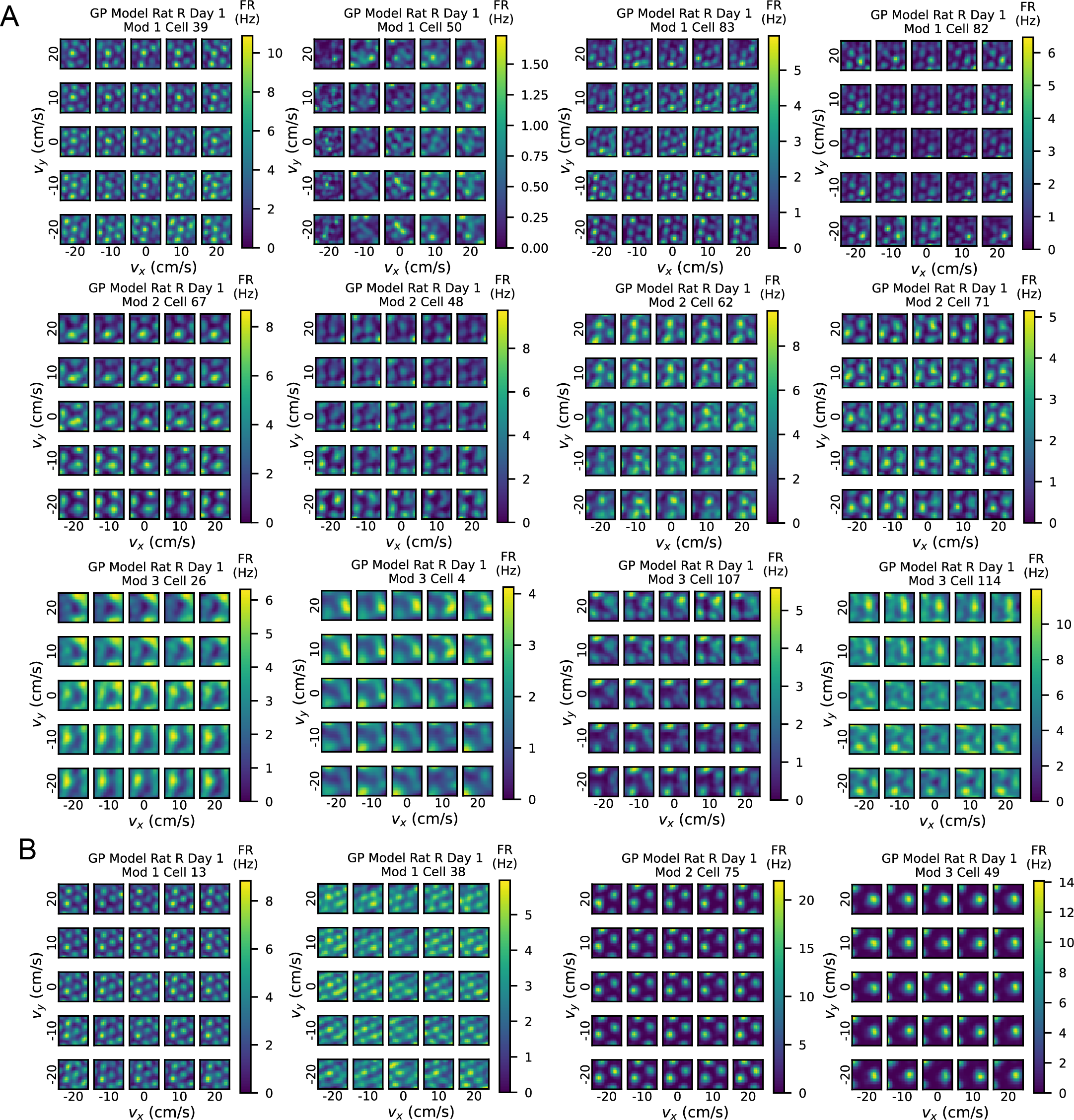

**Supplemental Figure 3: Additional examples of individual cells with non-separability in Rat R, Day 1.**

A) Cells with high SDCS (standard deviation of cosine similarity; see Methods) show non-separability in position-velocity tuning. Note that in Module 2, Cells 62 and 71 there is a positional shift in where the grid fields occur. Color bars indicate firing rate in Hertz as before.

B) Cells with low SDCS values exhibit similar tuning curves across velocity bins.

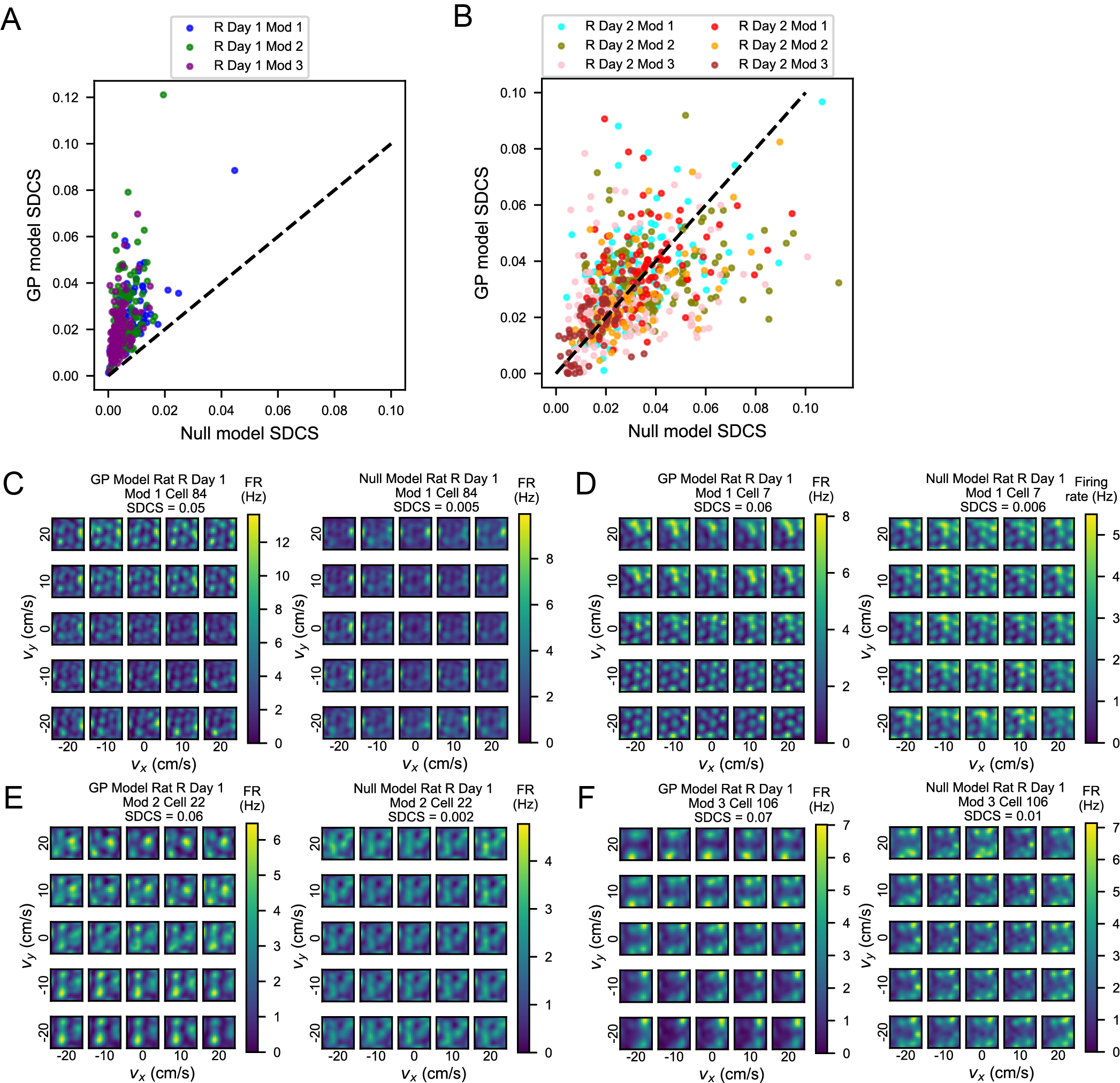

**Supplemental Figure 4:** **Analysis of GP performance on simulated data.**

A) Analysis of the validity of our GP model on sparse data. A null model was constructed by taking the position tuning curves of each cell at the 0 x- and y-velocity bin and simulating with a Poisson model the spike train at the same behavioral trajectories (see Methods). We compared the SDCS of the GP estimates of the 4D tuning curve for this null model to the SDCS of the GP estimates of the true data. In Rat R Day 1 Modules (1-3), all points lie above the unity line, meaning the non-separability we see in the data for this session cannot be explained only as an artifact of the GP process on sparse data.

B) The same analysis in A but shown on the other sessions (Rat R Day 2, Rat Q, and Rat S). SDCS values of the null model data are shown on the x-axis, and corresponding SDCS values of the real data are shown on the y-axis, each after being processed in our GP pipeline. On these sessions, the points lie scattered around the unity line, indicating that the data we encountered for these sessions was too sparse for the GP model to reliably extract signal.

C) For cell 84 from Rat R Day 1 Module 1, the left panel shows the results of running the GP model on the real data, while the right panel shows the same analysis on simulated data. Color bar shows the firing rate in Hz of the cell.

D) For cell 7 from Rat R Day 1 Module 1, the left panel shows the results of running the GP model on the real data, while the right panel shows the same analysis on simulated data. Color bar shows the firing rate in Hz of the cell.

E) For cell 22 from Rat R Day 1 Module 2, the left panel shows the results of running the GP model on the real data, while the right panel shows the same analysis on simulated data. Color bar shows the firing rate in Hz of the cell.

F) for cell 106 from Rat R Day 1 Module 3, the left panel shows the results of running the GP model on the real data, while the right panel shows the same analysis on simulated data. Color bar shows the firing rate in Hz of the cell.

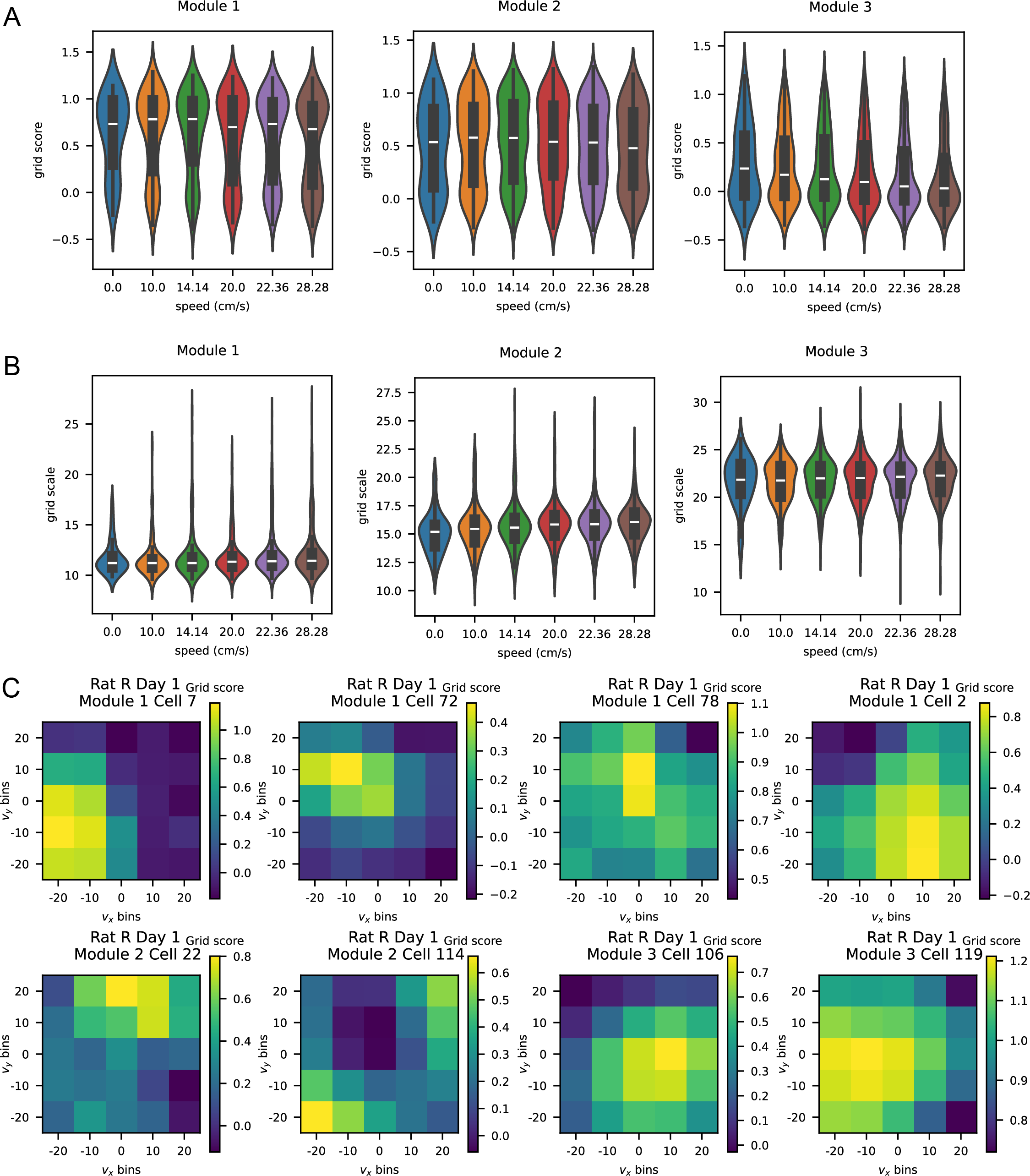

**Supplemental Figure 5: Additional characterization of non-separability on neurons within the top 50% of SDCS per module.**

A) Violin plot of the distribution of grid score across different speed bins. Aggregating velocity bins into speed bins by magnitude only shows that grid score does not change across speeds. Left panel shows grid scores in Rat R Day 1 Module 1, middle shows Module 2, and right shows Module 3.

B) Violin plot of grid scale values across different speed bins. The distribution of grid scale across different speed bins does not change. Left panel shows grid scores in Rat R Day 1 Module 1, middle shows Module 2, and right shows Module 3.

C) While there are no systematic differences in grid fields our analysis revealed, on an individual cell basis, some cells showed trends in grid score across velocity space. Heat maps show grid score across velocity space.

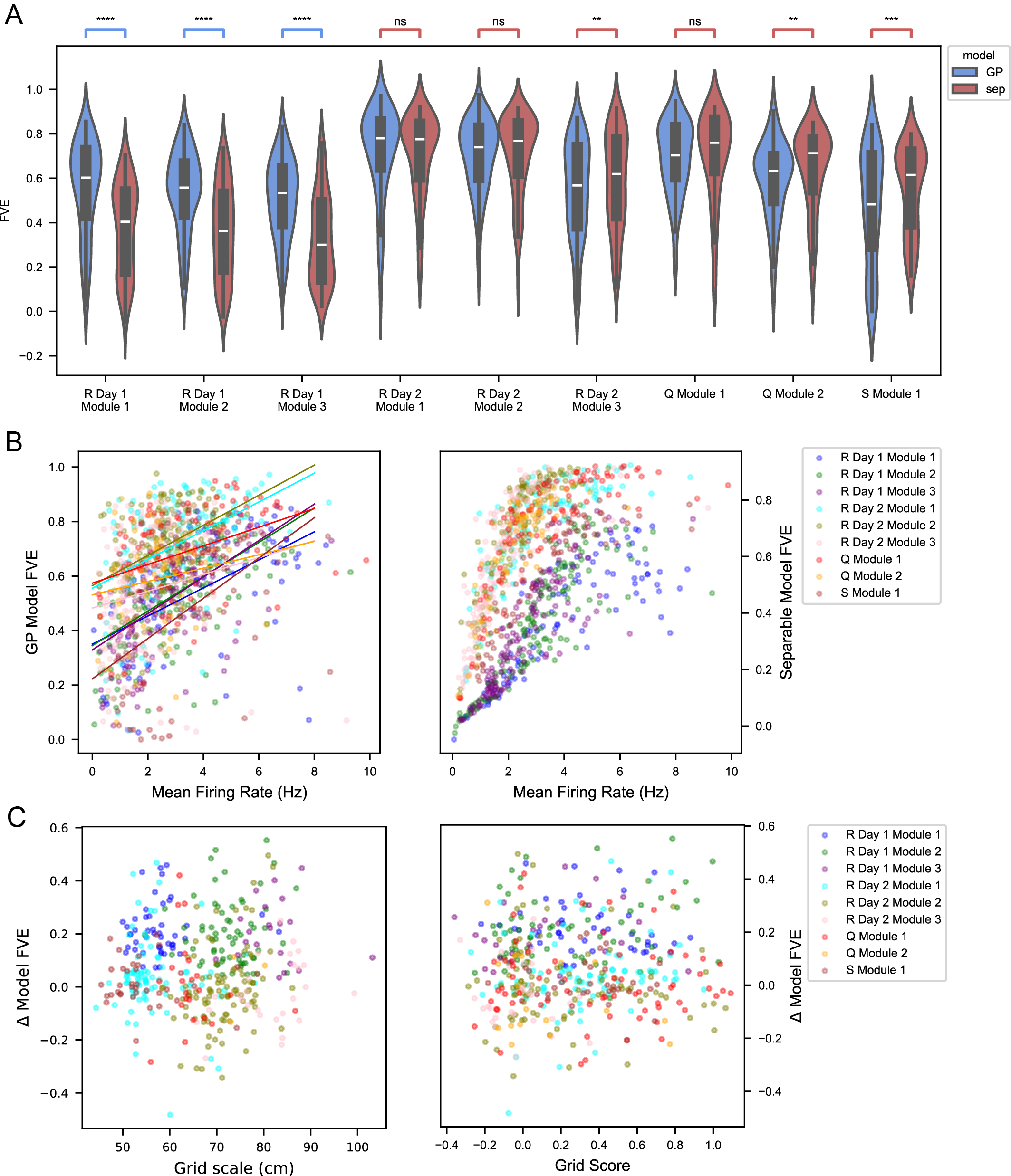

**Supplemental Figure 6: Differences in the performance of the GP and separable models are driven by mean firing rate but not grid score or grid scale.**

A) Violin plots showing the difference in FVE across models for each module and session. Significance bars are shown via Wilcoxon signed-rank test and are colored according to the direction of the test. Blue brackets indicate that the test was performed for the GP model FVE being greater than the separable model FVE, whereas red brackets indicate that the test was performed for higher separable model FVE. White line indicates the median, black boxes indicate the interquartile range (IQR), and black lines are drawn to 1.5 times the IQR or the most extreme point in the dataset.

B) Scatterplots showing mean firing rate on the x-axis and performance of the GP model (left) and separable model (right) on the y-axis. Each point is one cell, and sessions and modules are color-coded. GP model performance increases with mean firing rate. Legend on the right shows the color code for points in each session and module. C) Left: Scatterplot showing grid scale in cm on the x-axis with difference in model performance on the y-axis. Right: Scatterplot showing grid score on the x-axis and difference in model performance on the y-axis. Legend on the right shows the color code for points in each session and module.
